# Mechanism underlying the ultralow energy-consumption rapid ion dehydration for the high flux of KcsA potassium channels

**DOI:** 10.1101/2025.11.30.691452

**Authors:** Yue Wang, Bo Song, Lei Jiang

**Affiliations:** State Key Laboratory of Bioinspired Interfacial Materials Science, Suzhou Institute for Advanced Research, University of Science and Technology of China, Suzhou, Jiangsu 215123, China; Nano Science and Technology Institute, University of Science and Technology of China, Hefei 230026, China; Hubei Key Laboratory of Agricultural Bioinformatics, College of Informatics, Huazhong Agricultural University, Wuhan 430070, China; School of Optical-Electrical Computer Engineering, University of Shanghai for Science and Technology, Shanghai 200093, China; Institute for Biomedical Materials & Devices (IBMD), Faculty of Science, University of Technology Sydney, Sydney, NSW 2007, Australia

**Keywords:** resonance, tunneling-like ion dehydration, high flux, biological channels

## Abstract

High-flux and ultralow energy consumption (UEC) transport of biological and artificial ion channels has been widely reported [1-22]. However, there is a precondition for such transport: UEC rapid ion dehydration, which underlying mechanism is a remaining challenge. Here, by molecular dynamics (MD) simulations, we demonstrate that a K^+^ ion can transfer from the hydration water outside KcsA channel to the bound water in the channel without its hydration water accompanying, i.e., a tunneling-like motion, which provides a basis for the UEC dehydration of ions. In our simulations, a KcsA channel was divided into three regions: Cavity-1, Cavity-2 and the filter. As a K^+^ ion moves from Cavity-1 to Cavity-2, there occurs a resonant energy transfer to the ion from the filter-confined coherently oscillating ions, leading to a tunneling-like motion of the ion from the Cavity-1 to Cavity-2 water with no hydration shell accompanying and no influence on the Cavity-2 water. As the K^+^ ion further moves from Cavity-2 to the filter, the ion adjusts its hydration-shell water structure to coherence-resonantly couples with the filter-confined ions, leading to another tunneling-like motion to reach complete dehydration. These two consecutive processes constitute a "resonant tunneling” of the ion from Cavity-1 to the filter, as a basis for the UEC rapid dehydration. Our findings provide an understanding of the dehydration dynamics in biological channels and its relationship with the UEC high-flux, potentially promoting the development of artificial membranes design.

---

The high flux as well as the high selectivity is the character of biological ion channels, enabling the ultralow energy consumption (UEC) and ultrahigh accuracy of biological systems [1]. Inspired by the biological channels, a large amount of artificial membranes with nanopores are designed to achieve ion transport with a high flux (10^7^ ∼ 10^8^ ions·s^-1^) as well as high selectivity [2-21]. These advances in turn urgently call for the understanding of biological channels. Mackinnon et al. deeply studied ion selectivity of potassium channels a quarter century ago, and attributed it to the special structure of the ion-selective filter [22]. De Groot et al. proposed a direct Coulomb knock-on model for K^+^ ions permeating the channel, i.e., the Coulomb repulsion between adjacent ions is a key to the high-efficiency ion conduction [23]. Zanni et al. suggested a knock-on model in which K^+^ ions and water molecules exhibit alternate passage through the filter [24]. The further investigation of de Groot et al. showed that the completely-dehydrated K^+^ ions occupy all the filter sites in the transport process, i.e., a direct knock-on model [25]. Recently, we demonstrated a quantum coherence model for the high-ordered ion transport, i.e., the filter-confined K^+^ ions move together as a coherent body, which determines the high flux and UEC of KcsA channel [26]. Despite these advancements, there remains an unclear issue: As a precondition for the UEC high-flux transport of channels, a hydrated ion should conduct UEC rapid dehydration before it transports in the channel.

To achieve UEC ion dehydration in the channels, we believe that a pre-condition should be met, i.e., the hydration water of an ion does not accompany the ion into the channel; otherwise the reverse flow of ionic hydration water will result in massive energy consumption. Therefore, during the process, the number of channel-bound water molecules will remain unchanged. Applying the KcsA potassium channel as a typical example, we show this hypothesis detailedly. There are a hydrophilic gate and a hydrophobic gate in the channel (Fig. 1a). The channel is thus divided into three regions: a cavity (labeled Cavity-1) outside of the hydrophilic gate, a cavity (labeled Cavity-2) between the hydrophilic and hydrophobic gates, and a Filter+ area inside of the hydrophobic gate. A hydrated K^+^ ion initially outside the channel moves to the channel, and has no interaction with the channel-confined ions, while the ion will cause a change in the bound water of Cavity-1 as it is close to the channel, leading to an unoccupied area of water molecules for the ion, i.e., K^+^ hole_1_ (Fig. 1a t1, with the simulation results in the right panel). As the ion further moves close to the hydrophobic gate, an interaction with the channel-confined ions may form, while the bound water of Cavity-2 also changes, leading to an unoccupied area of water molecules for the ion, i.e., K^+^ hole_2_ (Fig. 1a t2, with the simulation results in the right panel). All these effects give a rise to the possibility that when a hydrated ion passes through the hydrophilic gate, the ion undresses its hydration shell and then redresses a new one in Cavity-2, i.e., a tunneling-like ion motion without its hydration water accompanying. Finally, passing through the hydrophobic gate and moving into Filter+, the ion undresses its hydration shell and achieves the complete dehydration (Fig. 1a t3). Therefore, no shell water accompanies the ion in the entire process of moving through the channel, and then the influence of ion dehydration on the bound H_2_O numbers in Cavity-1 and Cavity-2 should be ignorable.

**Fig. 1.**
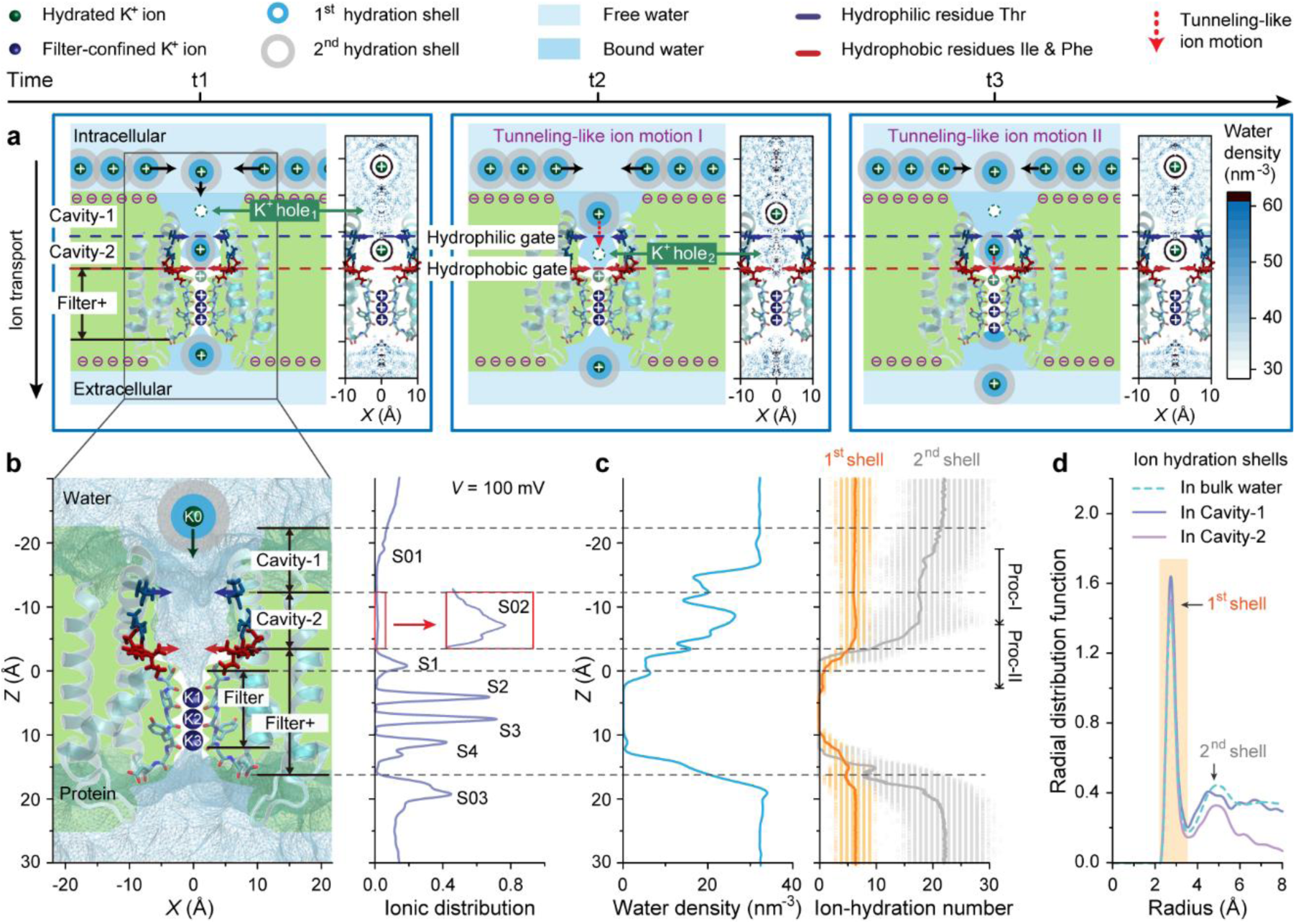
Hypothesis of tunneling-like K^+^ ion dehydration in a KcsA channel together with the molecular dynamics simulation system and preliminary analyses. **a)** Hypothesis of tunneling-like K^+^ ion dehydration in a KcsA channel. At the time t1, a hydrated ion outside of the channel moves to the channel Cavity-1, with the simulation results in the right panel. At t2 and t3, an ion conducts a tunneling-like motion (dotted red arrow) from the Cavity-1 to Cavity-2 water and from the Cavity-2 water to Filter+, respectively. The ion passes through the hydrophilic (dark blue) and hydrophobic (dark red) gates of channel without its hydration shell accompanying, different from the conventional diffusion of a hydrated ion. The water in Cavity-2 is unchanged in the processes, i.e., its changes caused by the ion is ignorable. **b-d)** KcsA channel system and water analyses of K^+^ ion dehydration. **b)** Simulation system (left) and ion-occupation sites (right) of KcsA channel under a transmembrane voltage (*V*) of 100 mV. The light blue network indicates the spatial distribution of water; the white and green parts denote the areas of protein pore and protein, respectively. **c)** Distributions of water density and ion hydration number. The orange and gray curves represent the ion 1^st^- and 2^nd^-shell water, respectively. Proc-I and Proc-II mean the dehydration-related processes of an ion moving from Cavity-1 to Cavity-2 and from Cavity-2 to the filter, respectively. **d)** Radial distributions of ion shell water in Cavity-1 (light blue) and Cavity-2 (violet).

To examine this hypothesis, employing molecular dynamics (MD) simulations on a KcsA channel system, we demonstrate that a K^+^ ion can transfer from the hydration water outside the channel to the bound water in the channel without its hydration water accompanying, i.e., a tunneling-like motion, as provides a basis for the UEC ion dehydration (Fig. 1b). During the process, the number of water molecules bound in the channel remains unchanged, while a K^+^ hole appears in Cavity-1 as a K^+^ ion is close to the channel, and a K^+^ hole appears in Cavity-2 as a K^+^ ion is in Cavity-1. Specifically, as a K^+^ ion moves from Cavity-1 to Cavity-2, there occurs a resonant energy transfer to the ion from the filter-confined coherently oscillating ions, causing a tunneling-like motion of the ion from the Cavity-1 to Cavity-2 water with no hydration shell accompanying and no influence on the H_2_O number of Cavity-2. As the K^+^ ion further moves from Cavity-2 to KcsA filter, the ion adjusts its hydration-shell water structure to coherence-resonantly couples with the filter-confined ions, leading to another tunneling-like motion of the ion to reach complete dehydration. Such two processes cause rapid ion dehydration, providing a basis for the UEC high-flux of channel.

## Results and Discussion

To conduct fine investigation on the dehydration process of a K^+^ ion when it passes through a potassium channel, we performed MD simulations on a KcsA channel system. The transmembrane voltage (*V*) of a neural K^+^ current is usually in the range of ∼80 mV [27]. An open KcsA channel system under *V* = 100 mV was thus applied (Fig. 1b left, Supplementary Fig. 1). There were seven sites for ion occupying in the channel, labeled S01, S02, S1, S2, S3, S4 and S03 (Fig. 1b right). A hydrophilic residue-formed gate was located between S01 and S02, and a hydrophobic residue-formed gate was between S02 and S1, labeled hydrophilic and hydrophobic gates, respectively. We thus divided the channel into three regions: Cavity-1, Cavity-2 and Filter+. The density distribution of water was calculated. As shown in Fig. 1c left, the density of water in the KcsA entrance terminal was 32.2 ± 0.3 nm^-3^, consistent with that (∼32 nm^-3^) of bulk water. A clear region of density minima between Cavity-1 and Cavity-2 was observed, due to the influence of the hydrophilic gate. In Cavity-2, there was a density maximum, related to the hydrophilic effect of threonine (Thr) residues on the Cavity-2 wall at *Z* ∼ -7.1 Å. From the Thr residues to filter, the density decreased, due to the influence of the hydrophobic gate. To explore the effects of the above two gates on the ionic shell water, we calculated the hydration number when the ion transports through the channel. The H_2_O number of 1^st^-shell water kept on a plateau of 6.4 ± 0.1 both in Cavity-1 and Cavity-2, and that of 2^nd^-shell water decreased from 20.2 to 17.2 in Cavity-1 and decreased to 9.5 from a plateau of 17.2 ± 0.2 in Cavity-2 (Fig. 1c right). The fluctuation of 2^nd^-shell H_2_O number was clearly limited when the ion was at the hydrophilic gate, meaning a clear influence of the gate on the hydration shell. In-between the hydrophobic gate and filter, the H_2_O numbers of ionic 1^st^ and 2^nd^ hydration shells decreased to 0.8 and vanishing, respectively, denoting complete dehydration. Although the radial distributions of 1^st^-shell water in Cavity-1 and Cavity-2 were very similar, those of 2^nd^-shell water were different, with maxima at 4.5 Å and 5.0 Å, respectively (Fig. 1d). Therefore, the water in Cavity-2 is clearly different from that in the other regions, while the significant changes in K⁺ ion hydration shells occur when the ion moves from Cavity-1 to Cavity-2 (labeled Proc-I) and from Cavity-2 to filter (labeled Proc-II), with the great effects of the hydrophilic and hydrophobic gates, respectively.

### Resonant energy transfer-driven tunneling-like dehydration when a hydrated K^+^ ion moves from the KcsA Cavity-1 to Cavity-2 (i.e., Proc-I)

To study the dehydration in Proc-I, we analyzed the structure change of water in Cavity-2 as a hydrated ion enters. The typical trajectories of hydrated ion *Z*-coordinate and Cavity-2 H_2_O number are shown in Fig. 2a left. For the time *t* = 350 ps to 380 ps, the hydrated ion K0 moved from *Z* = -14.3 Å to -8.5 Å, passing through the hydrophilic gate to Cavity-2. During the process, the number of H_2_O in Cavity-2 weakly fluctuated in a range of 33 ∼ 40 with an average of 36.5 ± 1.5, indicating that the change of Cavity-2 water is slight. The distribution of Cavity-2 H_2_O had a significant maximum at *Z* = -7.1 Å (the location of hydrophilic Thr), denoting that the Thr residues on the Cavity-2 wall have an attraction to the H_2_O molecules. Over all our simulations, the distributions of Cavity-2 H_2_O number in the absence and presence of K^+^ ion well matched each other, with an identical average value of 39 ± 3, meaning that the Cavity-2 H_2_O number is independent of the ion (Fig. 2a right upper). The previous error (±1.5) of Cavity-2 H_2_O number (36.5 ± 1.5) caused by K0 entering was in the range here (±3) of spontaneous Cavity-2 water fluctuation, indicating that the effect of K^+^ ion entry on Cavity-2 water is ignorable. All these analyses suggest that the hydration water accompanying the ion into Cavity-2 can be ignored, and the 39 ± 3 H_2_O molecules are well confined in Cavity-2 with help of hydrophilic Thr residues on the cavity wall. The property of Cavity-2 confined water over all the simulations was further studied. The number density of Cavity-2 water was 21.7 ± 1.7 nm^-3^ and 21.6 ± 1.7 nm^-3^ in the absence and presence of K^+^ ion, respectively, while that of bulk water was 32.15 ± 0.01 nm^-3^ (Fig. 2a right lower). The ion-caused slight change of only 0.5% in the Cavity-2 confined water density also supports that the shell water accompanying the ion into Cavity-2 is ignorable. The density of Cavity-2 water was significantly smaller than that of bulk water, with the fluctuation much larger than that of bulk water. Therefore, The water confined in Cavity-2 is in a state between liquid and gas, different from the incompressible liquid water, providing a feasibility for the tunneling-like motion of a naked ion from the Cavity-1 water to Cavity-2 water, different from the conventional diffusion of hydrated ions. Hence, we can conclude that the ion of a hydrated ion passes through the hydrophilic gate into Cavity-2 without the hydration shell accompanying (i.e., tunneling-like dehydration), and the ion-caused change in the Cavity-2 bound water is ignorable.

**Fig. 2.**
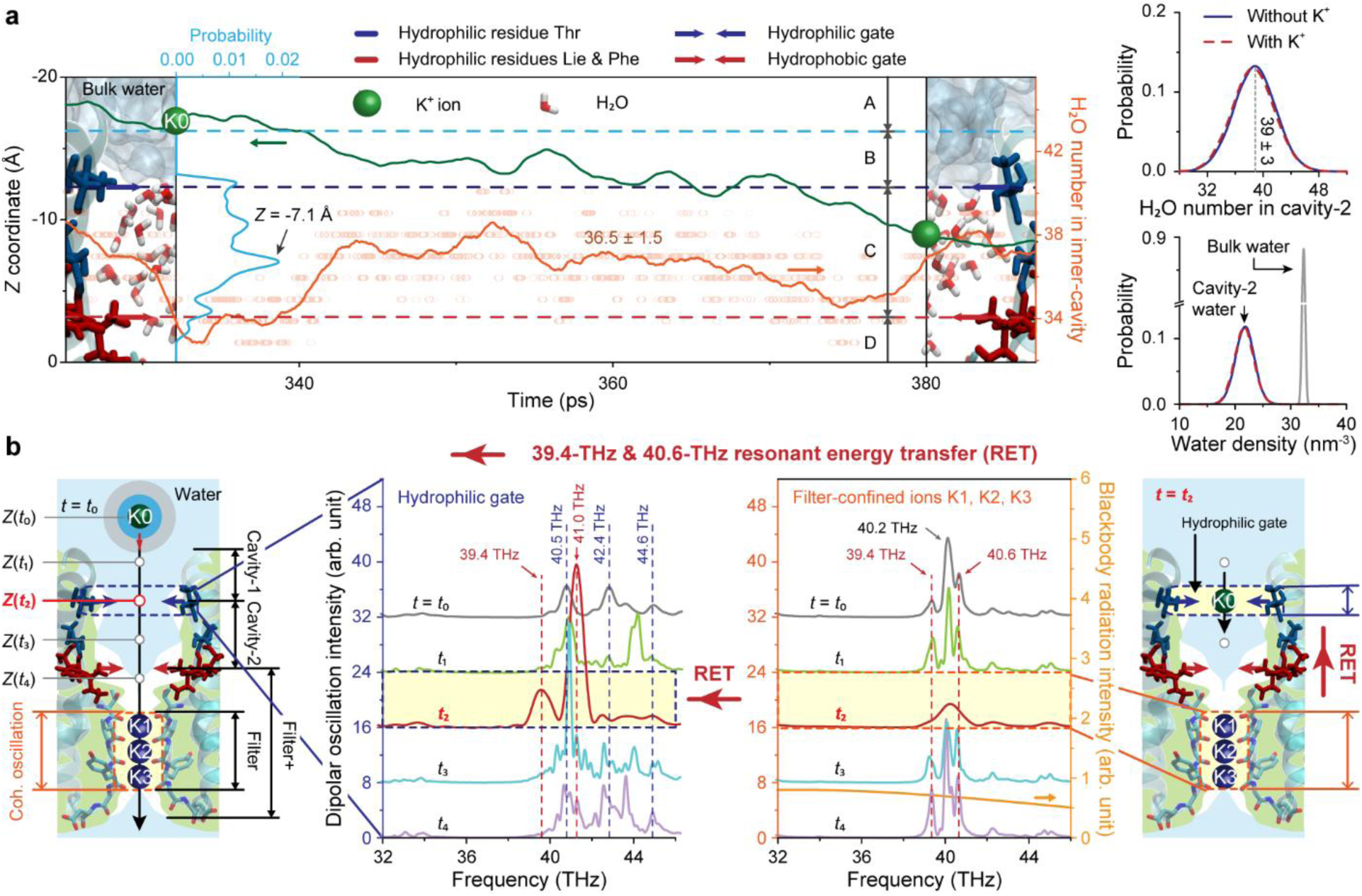
Resonant energy transfer-driven tunneling-like dehydration as a K^+^ ion moves from KcsA Cavity-1 to Cavity-2. **a)** Replacement of K^+^ ion hydration shell by a new one when the ion passes through the hydrophilic gate, i.e., tunneling-like ion dehydration. Left: Typical trajectories of the K^+^ ion coordinate (green) and Cavity-2 H_2_O number (tangerine, circles for the instantaneous values). The light blue curve stands for the H_2_O distribution in Cavity-2. The labels A, B, C and D means the areas out of the channel, in the channel and out of the hydrophilic gate, between the hydrophilic and hydrophobic gates, and in the hydrophobic gate, respectively. Right: Distributions of Cavity-2 H_2_O number (right upper) and water density (right lower) in absence (blue) and presence (red) of K^+^ ions over all the simulations. The gray curve indicates the bulk water. **b)** Resonant energy transfer (RET) from the filter-confined K^+^ ions (K1, K2, K3) to hydrated ion (K0) as the K0 passes through the hydrophilic gate. Left: Schematic representation of K0 moving from the outside of channel toward the filter. The label *Z*(*t_i_*) means the *Z* coordinate of K0 at the time *t_i_* (*i* = 0, 1, 2, 3, 4). Middle: Dynamics spectra of the hydrophilic gate (middle left) and filter-confined K^+^ ions (middle right). The gray, green, red, blue and violet curves denote the spectra at the time *t* = *t*_0_, *t*_1_, *t*_2_, *t*_3_ and *t*_4_, respectively. Right: Schematic representation of RET from the filter-confined ions to ion-coupled hydrophilic gate as the K0 is at the hydrophilic gate (*t* = *t*_2_).

To explore the driving force for a hydrated K^+^ ion to replace its hydration shell in the tunneling-like dehydration process, we analyzed dynamics spectra of KcsA ion transport. When a hydrated ion (K0) moved from the outside towards the filter, three ions (K1, K2, K3) were confined in the filter and performed coherent oscillation (Fig. 2b left) [26]. At the time *t* = *t*_0_, *t*_1_, *t*_2_, *t*_3_ and *t*_4_, K0 was successively outside the channel, in Cavity-1, at the hydrophilic gate, in Cavity-2, and in Filter+ (but outside the filter). Spectra of the filter-confined ion oscillation and hydrophilic gate vibration at the time *t_i_*, (*i* = 0, 1, 2, 3, 4) were calculated, respectively [26]. As shown in Fig. 2b middle, at *t* = *t*_0_, there were three vibration peaks of the hydrophilic gate at 40.5 THz, 42.4 THz and 44.6 THz, three oscillation peaks of the filter-confined ions at 39.4 THz, 40.2 THz and 40.6 THz, respectively. For *t* = *t*_1_ (K0 in Cavity-1), the changes in locations and intensities of the hydrophilic gate peaks clearly occurred, indicating that the K0 starts to couple with the gate, affecting the gate vibration. The peaks of filter-confined ions kept their initial states, denoting that the effect of K0 on K1, K2 and K3 is ignorable. At *t* = *t*_2_ (K0 at the hydrophilic gate), instead of the previous three peaks of hydrophilic gate, two huge peaks were observed at 39.4 THz and 41.0 THz, respectively, which means that the full coupling of K^+^ ion with the gate is achieved, resulting in a new vibration structure. As for the filter-confined ions, the 39.4-THz and 40.6-THz peaks disappeared, and the 40.2-THz peak intensity clearly decreased, denoting that the K0 at the hydrophilic gate has a significant long-range effect on the confined ions in the filter, causing a great energy loss of them. Remarkably, the disappearing 39.4-THz peak had an identical frequency to the first peak of ion-coupled gate, and the disappearing 40.6-THz peak well matched the second one (41.0 THz) in the frequency. These facts mean that two resonant channels between the filter-confined ions and hydrophilic gate occur at the time *t*_2_, and the oscillation energy of confined ions is resonantly transferred to the ion-coupled gate (Fig. 2b right). Huge peaks of the ion-coupled gate indicate huge amplitudes of the gate and hydrated ion vibration, providing a force for the ion to overcome its coupling with the shell water, as is followed by the ion redressing a new hydration shell in Cavity-2. During *t* = *t*_3_ to *t*_4_, both the spectra of hydrophilic gate and filter-confined ions recovered to their initial cases, meaning that the K0 effect disappears as K0 leaves. Therefore, when a K^+^ ion passes through the KcsA hydrophilic gate, there is resonant energy transfer from the filter-confined coherently-oscillating ions to the ion-coupled gate, which drives the ion to replace its hydration shell by a new one to achieve the tunneling-like dehydration.

Note: the blackbody radiation of surroundings at body temperature is located at the wavelength of ∼9 μm (∼34 THz in the frequency) with the full width at half maximum of 11 μm, well covering the filter-confined ion coherence oscillation of ∼40 THz (Fig. 2b middle right, bottom) [28]. In principle, after the ion leaves the hydrophilic gate, the spontaneous radiation of surroundings will supplement the resonantly-transferred energy of filter-confined ions with a high efficiency.

### Coherent resonance-caused tunneling-like dehydration when a hydrated K^+^ ion moves from the KcsA Cavity-2 to filter (i.e., Proc-II)

To study the dehydration in Proc-II, we analyzed the trajectories of ions and hydration number during a hydrated ion moves from Cavity-2 to the filter. Typical trajectories of the ion *Z*-coordinates (K0, K1, K2, K3) and the K0 1^st^-shell H_2_O number are shown in Fig. 3 top. Overall, in the time of *t* = 431.3 ns to 437.8 ns, as K0 oscillated and moved to the filter, its 1^st^-shell H_2_O number oscillated and decreased, indicating multiple adjustments of the hydration shell as the K0 moves. K1, K2 and K3 were confined in the sites S2, S3 and S4 of filter, respectively, with ignorable fluctuations in the trajectories from 431.3 ns to 432.05 ns and obvious fluctuations from 432.05 ns to 437.8 ns, i.e., K0 has an influence on filter-confined ions only as it is close to the hydrophobic gate. Specifically, in the time *t* = 431.3 ns to 432.05 ns, the K0 moved towards the hydrophobic gate, and its 1^st^-shell H_2_O number was on a plateau of 6.5 ± 0.4. For *t* = 432.05 ns to 435.8 ns, as the K0 oscillated and passed through the hydrophobic gate to site S1, the 1^st^-shell H_2_O number oscillated and declined from 6.3 to a plateau of 1.6 ± 0.1, meaning multiple adjustments of the hydration shell near this gate. In a short time from 435.8 ns to 436.1 ns (labeled the time region I), with K0 moving rapidly moving back to the hydrophobic gate, the shell water number significantly increased from 1.7 to 4.5, meaning a large adjustment of the hydration shell. For *t* = 436.1 ns to 437.8 ns, as K0 migrated from a plateau of *Z* = -2.2 ± 0.2 Å (close to the hydrophobic gate) to a plateau of *Z* = -0.4 ± 0.2 Å (close to the site S1), its shell H_2_O number decreased from plateaus of 4.0 ± 0.2 to 1.6 ± 0.1. After that, in a short time from 437.8 ns to 437.96 ns (labeled the time region II), K0, K1, K2 and K3 synchronously underwent rapid transfer, with K3 exiting the filter and the ions K0, K1, K2 moving to the sites S2, S3, S4, respectively. The K0 ion hydration number declined from 1.7 to 0.5, meaning complete dehydration. The water molecule of accompanying the K0 into the filter thus can be ignored, thereby the process of *t* = 432.05 ns to 437.96 ns can be regarded as tunneling-like dehydration too. Finally, in 437.96 ns to 439.0 ns, K3 left the channel, the K0, K1, K2 kept at the S2, S3, S4, respectively, and the K0 hydration number on a plateau of 0.5 ± 0.2. Thus, as the ion hydration shell structure is adjusted, another tunneling-like dehydration occurs when a K^+^ ion moves from Cavity-2 to the filter, while the ion has an influence on the filter-confined ions when it is close to the hydrophobic gate, potentially in turn playing a role in the ion dehydration.

**Fig. 3.**
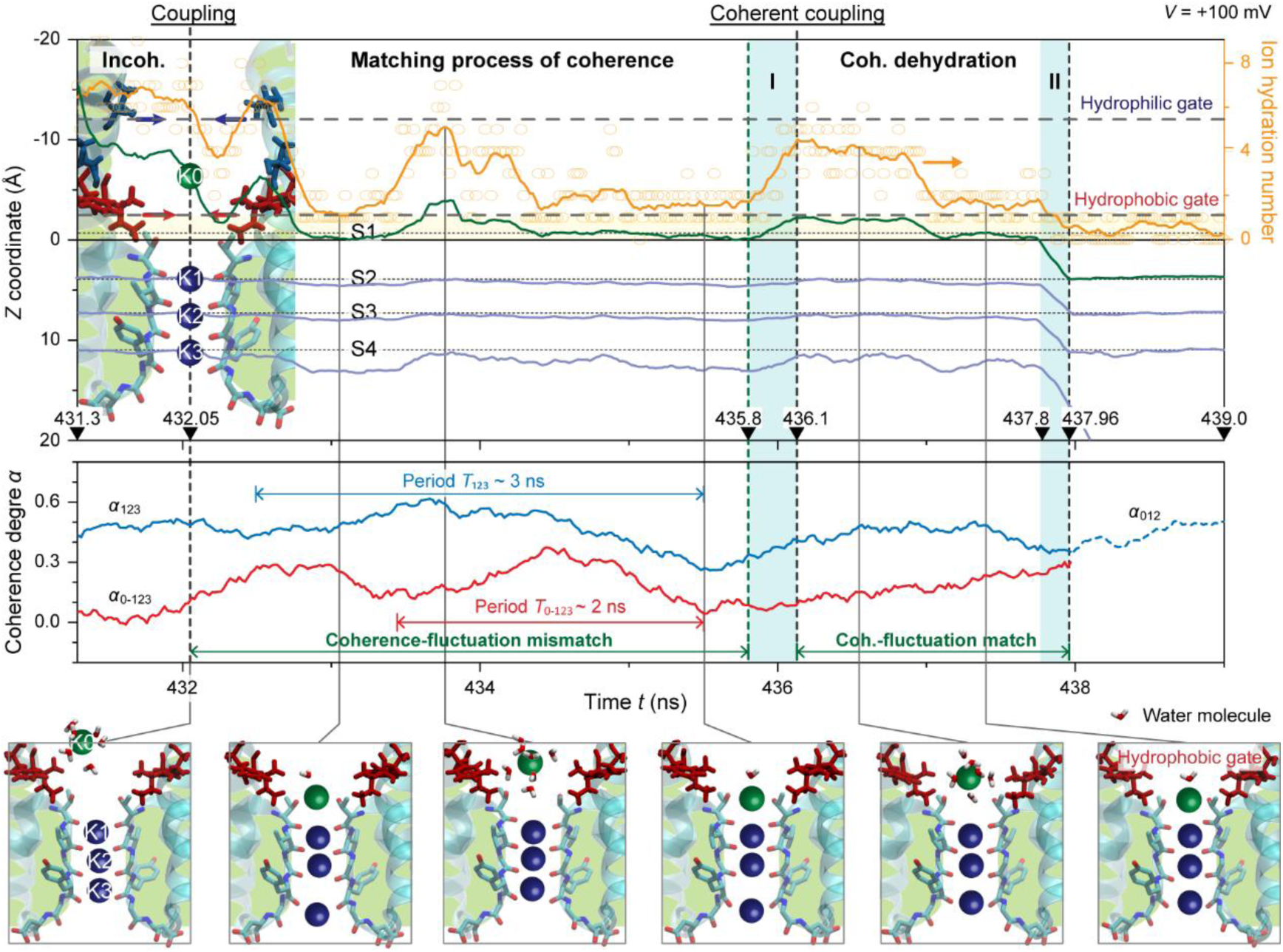
Coherent resonance-caused tunneling-like dehydration as a K^+^ ion moves from KcsA Cavity-2 to filter. The green and blue spheres indicate the K^+^ ions initially in the Cavity-2 (K0) and filter (K1, K2, K3), respectively. Top: *Z*-coordinate trajectories of K^+^ ions (green for K0 and gray blue for K1, K2, K3) and K0 1^st^-shell H_2_O number (orange, circles for the instantaneous values). Middle Trajectories of the ion coherence degree *α*. The red and light blue curves indicate the coherence degree of K0 with three filter-confined ions (K1, K2, K3) and that of the three filter-confined ions (blue solid: K1, K2, K3 for *t* < 437.96 ns, blue dashed: K0, K1, K2 for *t* > 437.96 ns), respectively. Bottom: Typical conformations of the hydrated K0 ion in the process.

To investigate the action of filter-confined ions on the ion of tunneling-like dehydration near the hydrophobic gate, we introduced the coherence degree of the hydrated ion with the filter-confined ions and that of the filter-confined ions. The coherence degrees were defined as,

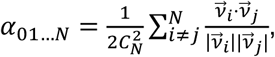

where the 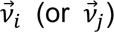 indicates the velocity of the *i*^th^ (or *j*^th^) K^+^ ion, with the ion labels *i*, *j* = 0, 1, 2, …, while *N* denotes the ion number. The coherence degree (*α*_0-123_) of the hydrated ion K0 with the filter-confined ions K1, K2 and K3 was calculated by the equation *α*_0-123_ = (*α*_01_ + *α*_02_ + *α*_03_)/3 with *N* = 2. The coherence degrees of three filter-confined ions, i.e., K1 – K3 (*i*, *j* = 1, 2, 3) for the time *t* = 431.3 ns to 437.96 ns and K0 – K2 (*i*, *j* = 0, 1, 2) for *t* = 437.96 ns to 439.0 ns, were calculated with *N* = 3, respectively, labeled *α*_123_ and *α*_012_.

Applying the coherence degrees above of ions, we analyzed the fine interaction process of a hydrated ion with the filter-confined ions as well as its effect on the tunneling-like dehydration. As shown in Fig. 3 middle and bottom, during *t* = 431.3 ns to 432.05 ns, the coherence degree *α*_123_ of filter-confined ions K1, K2 and K3 was on a plateau of 0.46 ± 0.02 with a very slight fluctuation, which corresponds to the fluctuation-ignorable ion coordinate trajectories of gray blue in Fig. 3 top. Thus, the K0 effect on the K1, K2 and K3 coherence is very weak when K0 is in Cavity-2, and the three filter-confined ions are in a coherent oscillation state. The coherence degree *α*_0-123_ of K0 with K1, K2 and K3 was less than 0.1, denoting that K0 is incoherent with other three ions, corresponding to the plateaus (Fig. 3 top, orange & green) of the hydrated ion. During *t* = 432.05 ns to 435.8 ns, the coherence degree *α*_123_ clearly fluctuated in the range of 0.26 to 0.61 with a period (*T*_123_) of ∼3 ns, different from the above plateau of *α*_123_. Namely, as K0 is close to the hydrophobic gate, assisted by the gate, a coupling of K0 with K1, K2 and K3 forms, causing a perturbation on the coherent oscillation of filter-confined ions. The degree *α*_0-123_ was in the range of 0.01 to 0.35 with a fluctuation period (*T*_0-123_) of ∼2 ns, meaning that the coupling of K0 with K1, K2 and K3 causes coherence between them. The difference of fluctuation periods *T*_123_ and *T*_0-123_ denotes that although the coherence occurs, the K0 still does not well coherently match the oscillation of the confined ions, i.e., they are in a coherence-matching process. In this process, to make K0 coherently match the filter-confined ion oscillation, the K0 shell water structure is multiply adjusted, i.e., the huge fluctuation (Fig. 3 top, orange) of hydration number. In the time region I of 435.8 ns to 436.1 ns, the degrees *α*_123_ and *α*_0-123_ increased from 0.31 to 0.40 and from 0.03 to 0.10, respectively. From 436.1 ns to 437.96 ns, *α*_123_ kept on a plateau of 0.44 ± 0.03, consistent with their initial plateau of 0.46 ± 0.02, while *α*_0-123_ increased from 0.10 to 0.30, and its difference with *α*_123_ (0.35) at 437.96 ns is small enough to ignore. Namely, by the last adjustment of hydration shell in the time region I, the coherent-resonance coupling of K0 with three filter-confined ions is achieved, causing a coherent body of four ions moving together towards the exit in the time region II (Fig. 3 top, green & gray blue). This process responds to that after the hydration-shell adjustment, the hydration number (Fig. 3 top, orange) of K0 ion decreases from 4.5 to 0.5. From 437.96 ns to 439.0 ns, as K3 left the channel, *α*_012_ kept on a plateau of 0.45 ± 0.02, consistent with the initial value (0.46 ± 0.02) of *α*_123_, meaning that the coherent oscillation of filter-confined ions recovers. Therefore, we can conclude that through adjusting the structure of K^+^ ion hydration shell assisted by the hydrophobic gate, the ion forms a coherent resonance with the filter-confined coherently-oscillating ions, causing the tunneling-like dehydration near this gate.

The consecutive two tunneling-like processes above of a K^+^ ion from Cavity-1 to Cavity-2 (i.e., Proc-I) and then to the filter (i.e., Proc-II) suggest a resonant tunneling from Cavity-1 to the filter. As shown in Fig. 4a, the hydrophilic and hydrophobic gates provide two barriers (labeled Barrier-1 and Barrier-2) in the ion pathway from the area A (outside of channel) to the areas B (Cavity-1 outside of the hydrophilic gate,), C (Cavity-2 between the hydrophilic and hydrophobic gates) and D (the filter inside of the hydrophobic gate). There are the bound states of ion at S01 and S02 in Cavity-1 and Cavity-2, respectively, labeled BS-1 and BS-2 (Fig. 4a lower), with a collective coherence oscillation state of ions confined in the filter. As indicated by our previous simulations (see Figs. 2,3), all the oscillation frequencies of BS-1, BS-2 and filter-confined ion coherence oscillation states are in the range of 39 ∼ 41 THz, i.e., they match each other, which enables a K^+^ ion resonantly tunneling from Cavity-1 to the filter through the bound state BS-2 between the two barriers.

**Fig. 4.**
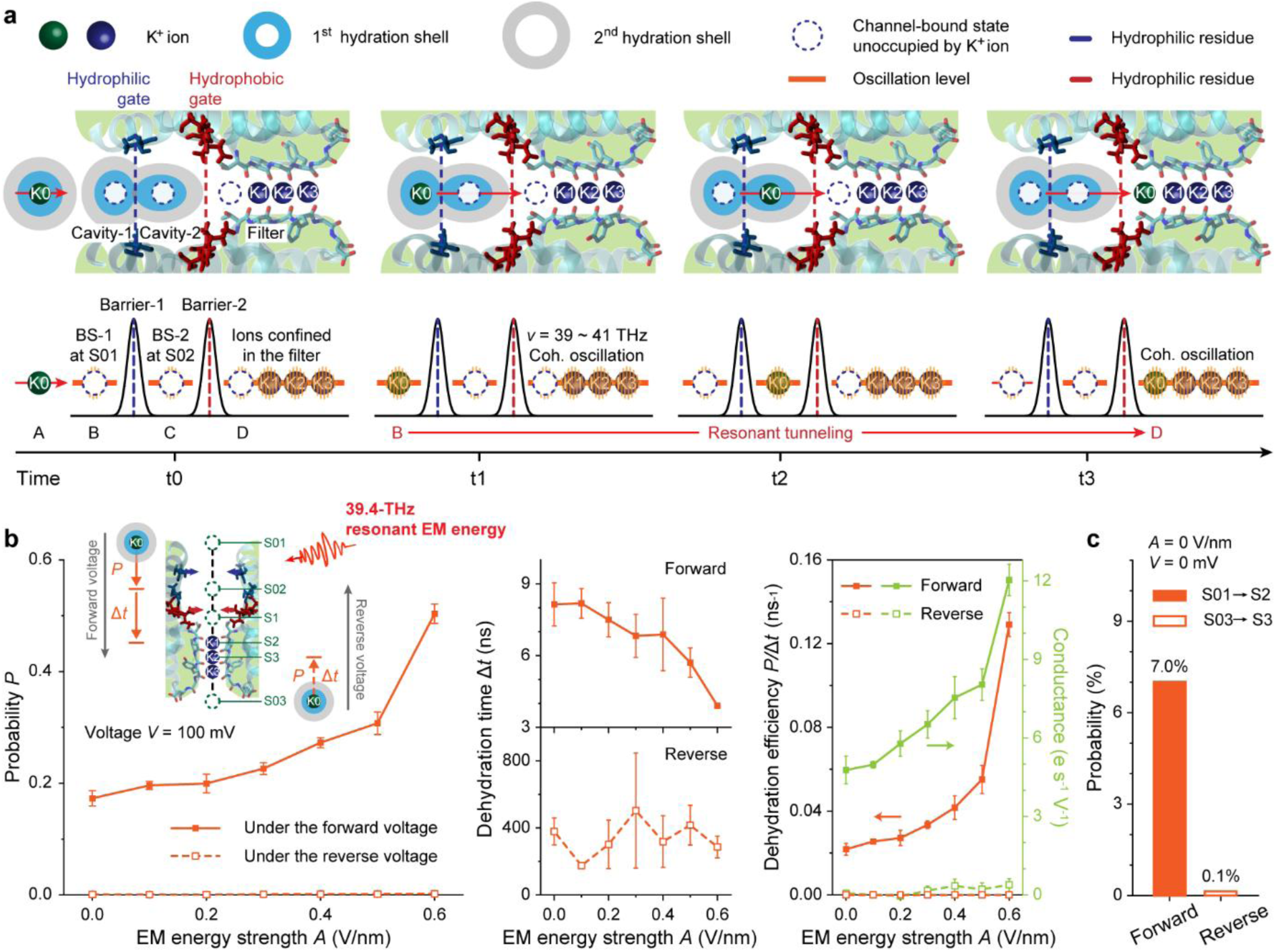
Resonant tunneling of ion from the KcsA outside to filter and its relation with the high flux of KcsA channel. **a)** Schematic representation of the resonant ion tunneling. The green and blue spheres indicate the K^+^ ions initially in the outside (K0) and filter (K1, K2, K3), respectively. Upper: Locations of K^+^ ions as the K0 moves from the KcsA outside to filter. The dark blue and dark red stands for the hydrophilic and hydrophobic gates, i.e., Gate-1 and Gate-2, respectively. Lower: Level structures for the K0 ion to perform the resonant tunneling. The labels A, B, C and D means the areas out of the channel, in Cavity-1, in Cavity-2, and inside of the hydrophobic gate, respectively. BS-1 and BS-2 indicate the bound state of ion at S01 and S02 in the areas B and C, respectively. **b)** Resonant energy effect on the dehydration and transport efficiencies of channel under the forward and reverse transmembrane voltages with an identical strength *V* = 100 mV. The resonant electro-magnetic (EM) energy, with a frequency (39.4 THz) identical to that of filter-confined ion coherence oscillation, is added to the channel (inset). Left: Probabilities (*P*) of the ion moving from the sites S01 to S02 (i.e., tunneling-like ion motion) under the forward voltage and from the sites S03 to S3 (i.e., conventional diffusion of hydrated ions) under the reverse, respectively. Middle: Time (Δ*t*) required for the complete dehydration as the ion moves from the sites S02 to S2 (tunneling-like motion) under the forward voltage and from the sites S03 to S3 (conventional diffusion) under the reverse one, respectively. Right: Dehydration efficiency (tangerine) and conductance (green) under the forward and reverse voltages. **c)** Effect of the resonant energy transfer of filter-confined ions on the forward and reverse ion motion in absence of the external EM energy and transmembrane voltage.

### Relationship of the tunneling-like ion dehydration with the high flux of KcsA channel

To study the effect of tunneling-like ion dehydration on the ion transport efficiency, we performed additional simulations with a frequency-fixed electromagnetic (EM) energy modulation, under the forward and reverse transmembrane voltages of 100 mV, respectively. As the previous analyses (see Fig. 2b), MIR 39.4 THz and 40.6 THz are the coherent oscillation frequencies of filter-confined ions, acting as the resonance channels of transferring energy for the tunneling-like dehydration. Therefore, a MIR EM field of 39.4 THz as a typical example was used to simulate the transferred energy from filter-confined ions, with the strength varied from 0.0 V/nm to 0.6 V/nm [26,29]. Under the forward voltage, the tunneling-like dehydration occurs when an ion passes through the hydrophilic gate to Cavity-2 (i.e., Proc-I) and moves from Cavity-2 to filter (i.e., Proc-II), respectively. To describe the dehydration in the processes, we applied the probability (*P*) of an ion passing through the hydrophilic gate, and the dehydration time (Δ*t*) of an ion moving from Cavity-2 to the site S2, respectively (Fig. 4b inset). Under the reverse voltage, an ion achieved complete dehydration as it moved from the sites S03 to S3 (Supplementary Fig. 2). The probability *P* and required time Δ*t* of this process were applied for the dehydration. Based on the *P* and Δ*t*, we introduced a parameter (*γ*) with an identical formula *γ* = *P*/Δ*t* for the dehydration efficiencies under forward and reverse voltages, respectively.

By these parameters of ion dehydration dynamics, we analyzed the relationship of tunneling-like ion dehydration with the high flux of KcsA channel. As shown in Fig. 4b, when the EM energy strength increased from 0.0 V·nm^-1^ to 0.6 V·nm^-1^, under the forward voltage, the probability *P* increased from 0.17 ± 0.01 to 0.50 ± 0.02, and the time Δ*t* decreased from 8.14 ± 0.90 ns to 3.91 ± 0.08 ns. The 39.4-THz resonant energy thus can promote both the ion capability of passing through the hydrophilic gate and the ion motion of Cavity-2 to the filter. Under the reverse voltage, the *P* and Δ*t* kept on plateaus of ∼10^-3^ and 339.9 ± 97.4 ns, respectively, denoting an ignorable effect of resonant energy on the process of a hydrated ion moving from the channel exit to filter (i.e., conventional diffusion of hydrated ions). The forward and reverse dehydration efficiencies *γ* exponentially increased from 0.022 ± 0.003 ns^-1^ to 0.129 ± 0.006 ns^-1^ and kept on a plateau of 7.8 ± 8.2 10^-6^ ns^-1^, respectively. The forward and reverse conductance of channel exponentially increased from 4.8 ± 0.5 e·s^-1^·V^-1^ to 12.0 ± 0.6 e·s^-1^·V^-1^ and only slightly shifted from plateaus of 6.6 ± 4.1 10^-4^ e·s^-1^·V^-1^ to 40.0 ± 7.3 10^-4^ e·s^-1^·V^-1^, respectively. Namely, the dehydration efficiency and conductance followed almost same laws under forward and reverse voltages, respectively, suggesting that the dehydration determines the transport. Remarkably, in the case of EM energy strength at 0.0 V·nm^-1^, i.e., no external EM energy, there is still much large differences in the ion dehydration efficiency and then conductance between the forward and reverse motions, which can be attributed to both the 39.4-THz and 40.6-THz energy resonantly transferred from the filter-confined coherently-oscillating ions. Therefore, the resonant energy transfer in the channel can efficiently promote the tunneling-like ion motion under a forward voltage, but cannot affect the conventional ion diffusion under a reverse voltage, causing the forward rapid-dehydration and then the forward high-flux of channel.

To further reveal the intrinsic difference between the forward and reverse fluxes of KcsA channel, we analyzed the ion dynamics under no transmembrane voltage *V* and no external EM energy. There still exists a 70:1 difference between the probabilities of K^+^ ion moving into the filter from the channel entrance and exit at *V* = 0 and *A* = 0 (Fig. 4c). This intrinsic asymmetry of channel can be attributed to the fact that even if there is no voltage, the resonant energy transfer from the filter to the ion-coupled hydrophilic gate still can occur, which provides an intrinsic advantage in the forward flow of hydrated ions, equivalent to a limitation on the reverse flow.

Finally, it is worth noting that the UEC rapid ion dehydration and transport highly depend on the ordered structure of ions confined in the channel. Additional electrical and mechanical stimulation will cause a structural change in the channel and a disorder in the channel-confined ions, leading to a dramatic decay in the efficiency of channel ion dehydration and transport. As an inference of our hypothesis, this phenomenon has been observed experimentally, i.e., an electromechanical coupling causes a structure change and consequent inactivation of the potassium channels [30,31].

In summary, by MD simulations on the KcsA channel system, we demonstrate that a K^+^ ion can transfer from the hydration water outside the channel to the water bound in the channel without its hydration water accompanying, i.e., a tunneling-like motion, which provides a basis for the UEC dehydration of ions. During the process, water in the channel remains unchanged. As a K^+^ ion moves from KcsA Cavity-1 to Cavity-2, there occurs a resonant energy transfer to the ion from the filter-confined coherently oscillating ions, causing a tunneling-like motion of the ion from the Cavity-1 to Cavity-2 water with no hydration shell accompanying and no influence on the Cavity-2 water. As the K^+^ ion further moves from Cavity-2 to the filter, the ion adjusts its hydration structure to coherence-resonantly couples with the filter-confined ions, leading to another tunneling-like ion motion to reach complete dehydration. These two consecutive processes constitute a “resonant tunneling” of the ion from Cavity-1 to the filter, providing a basis for the high flux of KcsA channel. Our findings improve the understanding of physics under the UEC and high flux of ion transport in life systems, potentially promoting the developments in treatment strategies for ion channel-associated diseases and health problems, and in design of bio-inspired UEC materials and devices.

## Methods

### Model preparation

We generated an open-state model of the KcsA channel employing the methodology outlined by Koper et al. Specifically, the high-resolution crystal structure (PDB ID: 1K4C) was used for the selectivity filter and the low-resolution open-state structure (PDB ID: 3F5W) was used for the remaining regions (Supplementary Fig. 1). This approach was necessitated by the dearth of high-resolution (≤ 2 Å) structures depicting the open-state KcsA potassium channel, despite the availability of more than 100 crystallographic entries. The PROPKA tool was used to predict the protein protonation state at pH 7.0. Modifications to the protonation state of Glu71 were carried out following Bernèche’s method. The membrane system was generated using CHARMM-GUI, in which the KcsA protein was embedded in a POPC lipid bilayer consisting of 216 POPC molecules. The system was solvated in a 9 × 9 × 10 nm^3^ simulation box containing 16,848 water molecules, 109 K^+^, and 109 Cl^−^ ions, corresponding to an ionic concentration of 0.4 M.

### Simulation setting

All molecular dynamics simulations were performed using GROMACS 2019, with the FF14SB and Slipids force fields used for the protein and lipid components, respectively. Water molecules were modeled using the TIP3P model, and ion parameters were taken from Lee et al. All bonds involving hydrogen atoms were constrained using the LINCS algorithm, allowing a time step of 2 fs, and the Verlet scheme was applied to update the neighbor list. Long-range electrostatic interactions were calculated using the particle mesh Ewald (PME) method, with a cutoff of 1.2 nm for electrostatic and van der Waals interactions. Temperature was maintained at 310 K using the V-rescale thermostat, and pressure was maintained at 1 bar using the Parrinello-Rahman barostat with semi-isotropic pressure coupling.

The single bilayer membrane system generated by CHARMM-GUI was energy-minimized and equilibrated for 20 ns with all heavy atoms except those in the selectivity filter (TVGYG) restrained with a force constant of 1000 kJ/mol/nm^2^. After equilibration, the system was duplicated along the z-axis to construct a composite bilayer membrane-protein system. The resulting system comprised 432 POPC lipids, 33,696 water molecules, 218 K^+^ ions, and 218 Cl^-^ ions in a 9 × 9 × 20 nm^3^ simulation box. The composite system was subsequently energy-minimized and equilibrated for an additional 20 ns under the same restraint conditions. Finally, production simulations were performed using the computational electrophysiology (COMPEL) protocol implemented in GROMACS for 1 µs under a fixed charge imbalance (Δ*q* = 4 e) across the membrane. The charge imbalance Δ*q* led to an ion concentration gradient Δ*C* of 14.6 mM and a resultant transmembrane voltage *V*. The gmx potential tool was used to calculate *V* using a relative dielectric constant (*ε*_r_) of 4. All statistical analyses were based on three independent 1-μs simulations.

## Acknowledgements

This work was supported by the National Key R&D Program of China (2021YFA1200404, B.S.), the National Natural Science Foundation of China (T22410002, B.S.; T2394532, B.S.).

## Author contributions

B.S. and L.J. conceived the study. B.S. designed the MD simulations, and Y.W. performed the MD simulations. All authors analyzed the results. B.S., L.J. and Y.W. wrote the manuscript.

## Supplementary information

Supplementary information is available on-line for this paper.

## Figures and Legends

**Supplementary Fig. 1.**
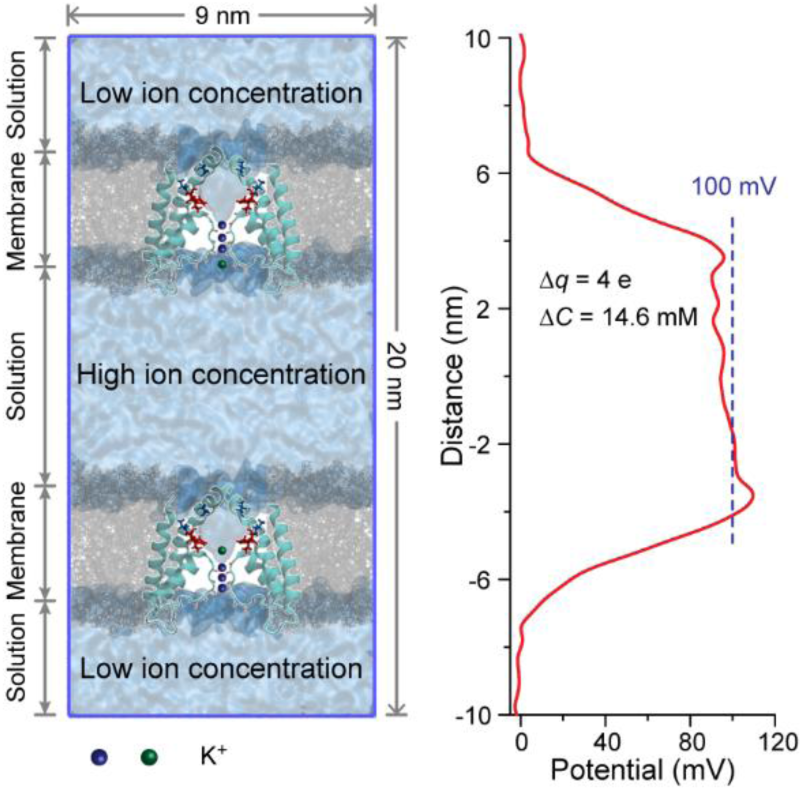
The whole simulation system and its potential change caused by transmembrane ion concentration difference. The labels Δ*q* and Δ*C* mean the charge imbalance and ion concentration difference, respectively.

**Supplementary Fig. 2.**
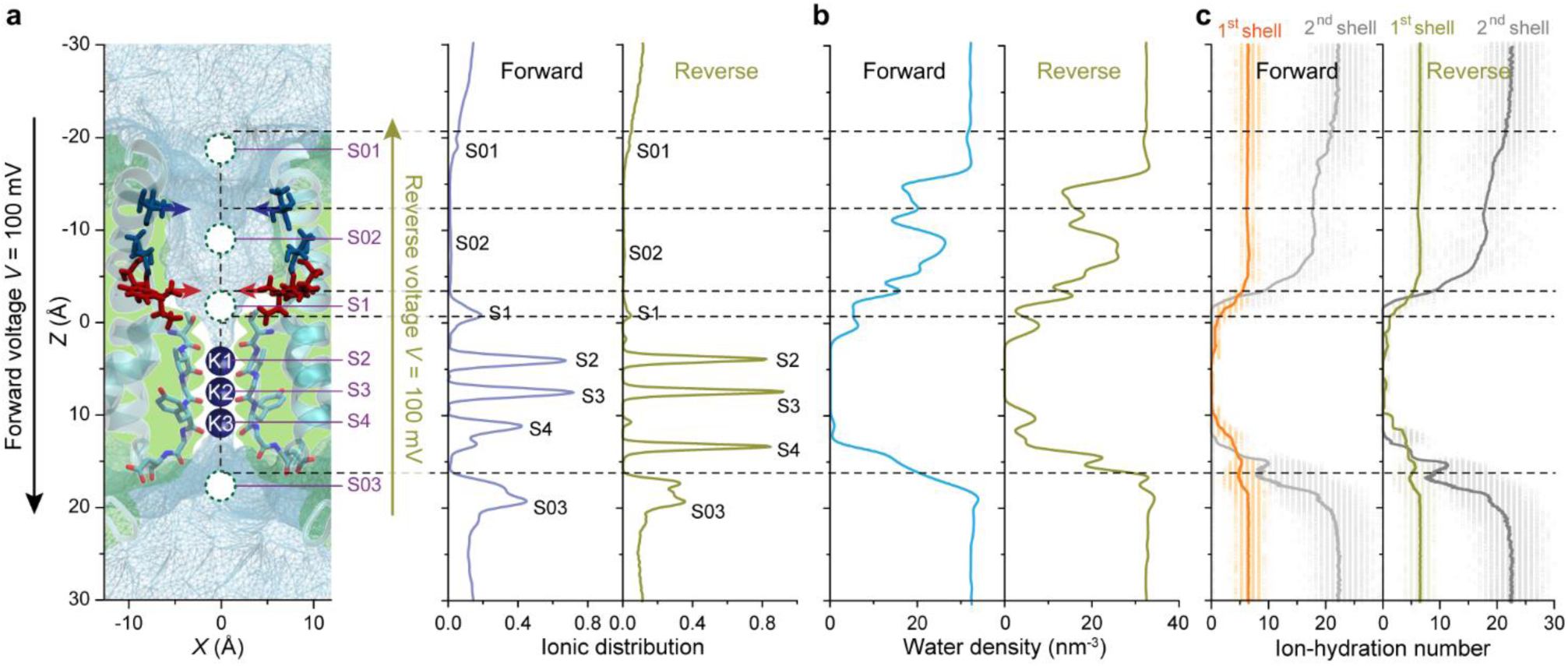
Distribution analyses of ions and water molecules in the KcsA channel under forward and reverse voltages. **a)** Ion-occupation sites in the channel under forward (middle) and reverse (right) transmembrane voltages (*V*) of 100 mV. **b)** Distributions of water densities under the forward (left) and reverse (right) voltages. **c)** Ion hydration numbers of the 1^st^- and 2^nd^-shell water under the forward (left) and reverse (right) voltages.

